# Global fingerprint of humans on the distribution of *Bartonella* bacteria in mammals

**DOI:** 10.1101/249276

**Authors:** Hannah K. Frank, Elizabeth A. Hadly

## Abstract

As humans alter habitats and move themselves and their commensal animals around the globe they change the disease risk for themselves, their commensal animals and wildlife. *Bartonella* bacteria are prevalent in many mammalian taxa, responsible for numerous human infections and presumed to be an important emerging group of zoonoses. Understanding how this genus has evolved and passed between host taxa in the past can reveal not only how current patterns were established but identify potential mechanisms for future cross-species transmission. We analyzed patterns of *Bartonella* transmission and likely sources of spillover using the largest collection of *Bartonella gltA* genotypes assembled, 860 unique genotypes of *Bartonella* globally. We provide support for the hypothesis that this pathogenic genus originated as an environmental bacterium before becoming an insect commensal and finally vertebrate pathogen. We show that rodents and domestic animals serve as the reservoirs or at least key proximate host for most *Bartonella* genotypes in humans. We also find evidence of exchange of *Bartonella* between domestic animals and wildlife and between domestic animals, likely due to increased contact between all groups. *Bartonella* is a useful infection for tracing potential zoonoses and demonstrates another major impact of humans on the planet. Care should be taken to avoid contact between humans, domestic animals and wildlife to protect the health of all.

## Introduction

Human movements and actions have numerous impacts for wildlife disease (1, 2). These impacts are of concern both from a wildlife conservation standpoint (1, 2) and from a public health perspective (spillover). Over 60% of emerging infectious diseases in the world are zoonotic, meaning they are transmitted from animals to humans (3). Despite the fact that zoonosis is an important component of emerging infectious diseases, it is often difficult to trace the ecology and evolution of zoonotic pathogens (4). Most efforts to identify the source of zoonoses occur after a human has become infected. Because spillover events are rare and often infection prevalence in the reservoir species is low, it can be difficult to trace the origin of potential zoonoses. However, *Bartonella* bacteria are an exception to this pattern. This genus of bacteria has been found in numerous taxa and is usually at high prevalence (5). *Bartonella* is a blood-borne pathogen, found in many animals. It is the cause of cat scratch fever, Carrion’s disease and trench fever as well as a number of incidents of endocarditis in humans and has been hypothesized to be the cause of unexplained febrile illness in a number of cases (5, 6). Therefore, it is an ideal pathogen to focus on in tracing zoonotic potential as well as potential impacts on the native hosts. In this study, we construct some of the largest global phylogenies to date of *Bartonella* from both 16s rRNA genes and citrate synthase (gltA) to determine the evolutionary history of *Bartonella*, patterns of host switching and geographic constraint and its spillover into humans or from human commensals into wild species. The citrate synthase gene is known to give high power to discriminate between *Bartonella* strains and is one of the most commonly sequenced *Bartonella* genes (7, 8). We also examine 16S as it is the most commonly sequenced locus for metagenomics studies, though it gives low power to discriminate between *Bartonella* species (8).

## Results

Starting with 1,618 *gltA* records, we analyzed 860 unique 277 bp sequences. In our final dataset, the most commonly sampled taxa were rodents (N = 559) and bats (N = 204), though some genotypes were found in multiple taxa so these are not strict numbers. Many nodes did not have good support so we conducted all analyses using only nodes with a posterior value of 0.7 or above. In order to test hypotheses regarding the timing of *Bartonella* cross-species transmission we created a time calibrated phylogeny using a subset of 334 sequences from which we were able to obtain a 548bp fragment of *gltA* and calibrated it at two nodes using the divergence time between hosts as the divergence time between *Bartonella* genotypes (see Materials and Methods). For this we used Costa Rican bats to calibrate the tree as their phylogenetic relationships have been well studied and there is no evidence to suggest *Bartonella* host shifts in this clade have been impacted by humans.

### Evolutionary history

Phylogenetic hypotheses generated from a 277bp fragment of *gltA* and a 259 bp fragment of the 16s rRNA gene both support an origin for *Bartonella* in the environment and in the guts of insects (both ectoparasitic and non-ectoparasitic species; Figures 1 and 2). Twice *Bartonella* has infected mammals from these environmental samples, which are basal to the main clade of mammal-associated *Bartonella* (Figure 2), which likely invaded mammals approximately 56 million years ago based on our time calibrated phylogeny, though it is unclear which mammalian host is ancestral (Figure 2; Figure S1).

**Figure 1:**
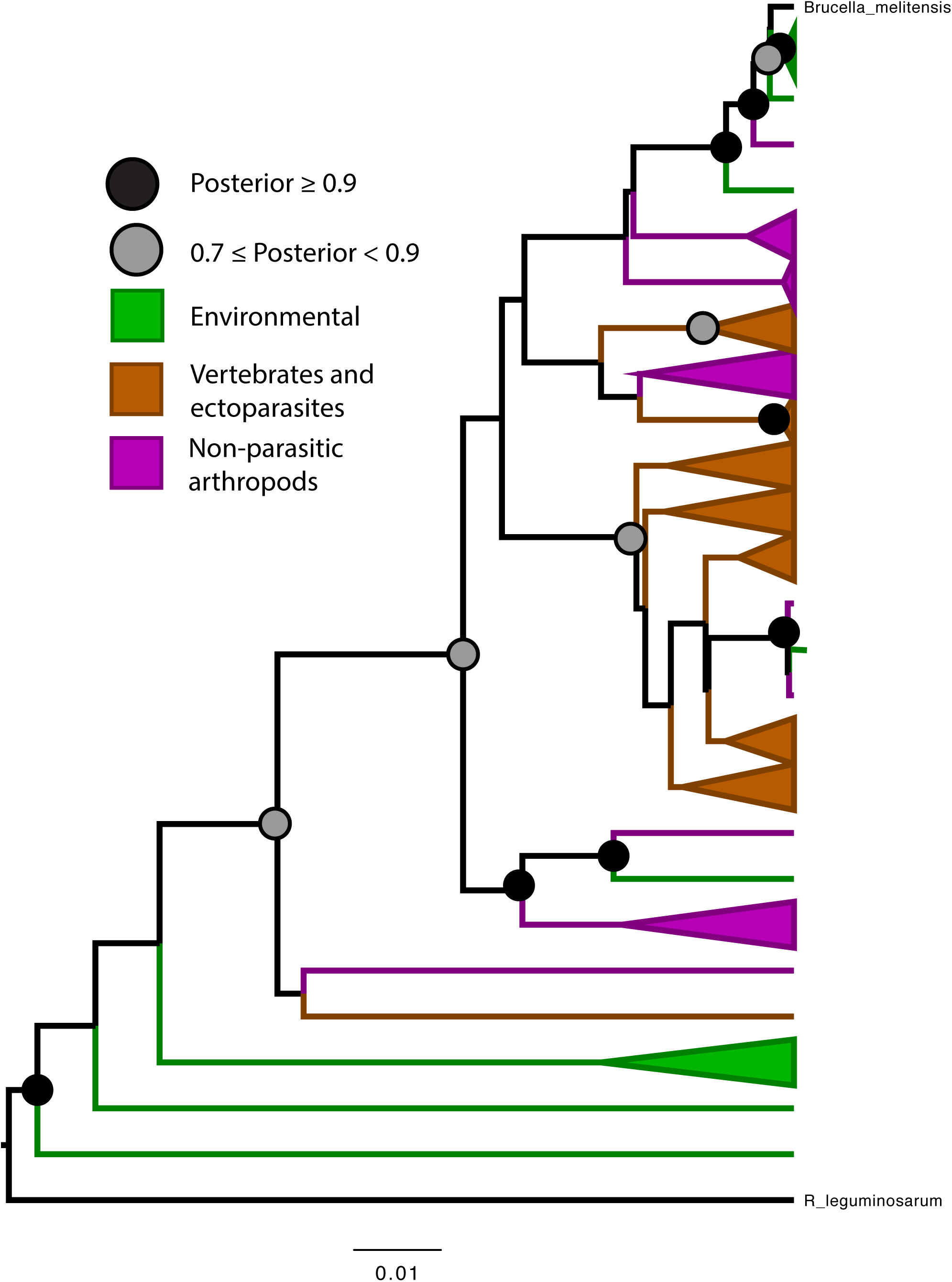
Bayesian phylogenetic hypothesis of *Bartonella* genotypes based on a 259bp fragment of 16s rRNA gene. Ectoparasites and their vertebrate hosts are colored brown; environmental sequences are green; non-ectoparasitic arthropods are colored purple. Scale bar indicates substitutions per site.

**Figure 2:**
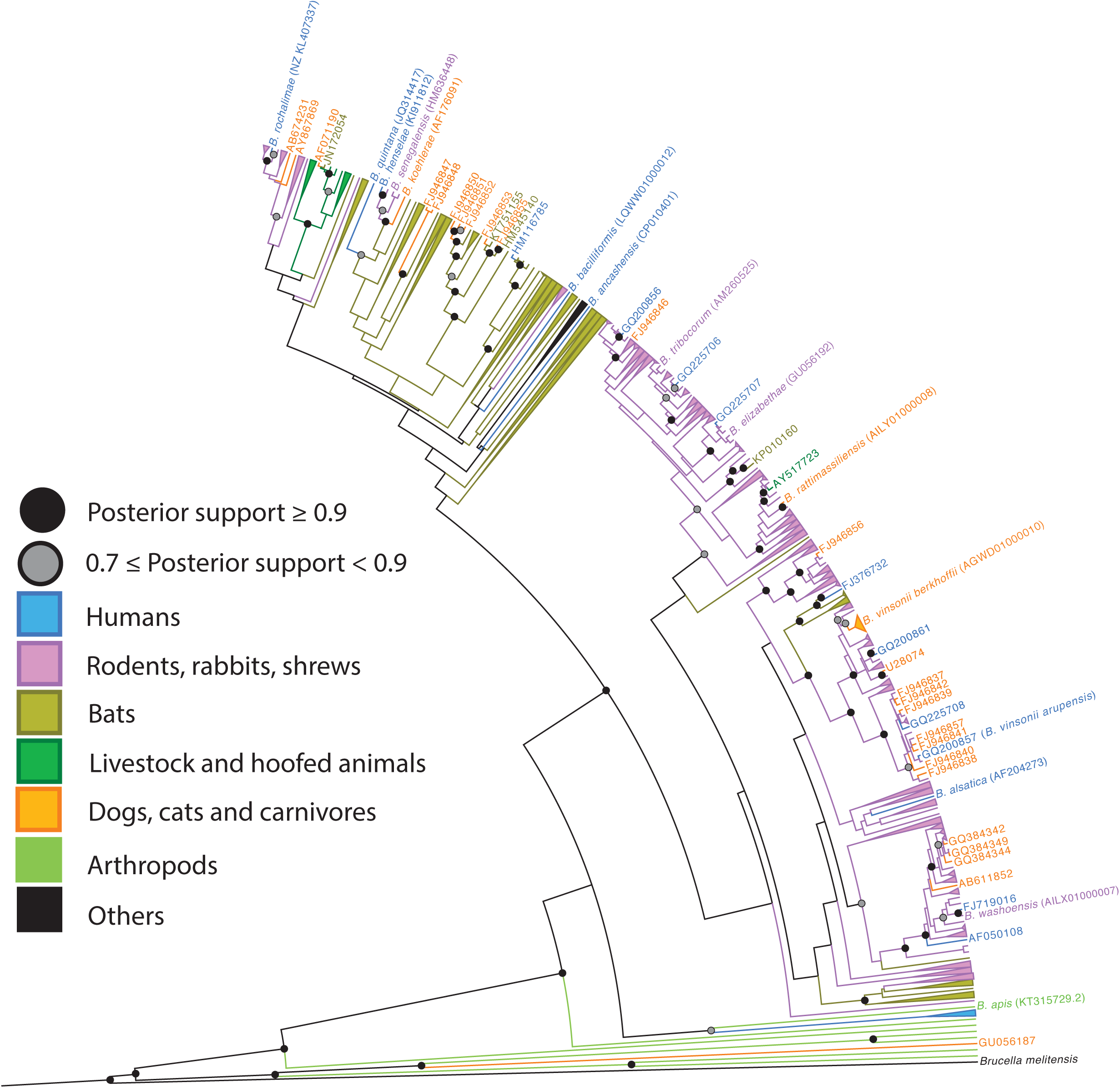
Bayesian phylogenetic hypothesis of *Bartonella* genotypes based on a 277 bp fragment *gltA*. Tip labels and branches have been colored according to the taxa in which they were identified with ectoparasites colored according to their host and collapsed to highlight specific patterns.

### Host and Geographic Conservation

*Bartonella* are generally highly host specific with closely related genotypes found in the same order of host; the best model of evolution for *Bartonella* host order in the 860 analyzed 277bp fragments was a lambda model in which lambda was 0.98, indicating near Brownian motion evolution along the phylogeny (AICc weight = 0.998; Table S1). Similarly, closely related genotypes of *Bartonella* were generally in the same geographic regions, whether analyzed by the continent from which the genotype was isolated or Old World versus New World (best model for both was a lambda transformation; continent: λ = 0.99, AICc weight = 0.88; OW-NW: λ= 0.99; AICc weight = 0.92; Tables S2 and S3).

### Exceptions to host specificity and limited geographic range: zoonosis and the human-domestic-wildlife interface

Despite the overall high host specificity of *Bartonella*, we observed a number of host shifts in our large phylogenetic hypothesis. Of 18 spillovers into humans (Figure 1; Table S4), 10 were from rodent clades, though two of these ‐‐ *B. clarridgeiae* and *B. henselae* – usually infect humans from domestic dogs and cats (9). In some cases, multiple strains of the same *Bartonella* species have infected humans representing separate spillovers, such as in the case of *B. washoensis* or the *B. vinsonii* complex which is found in both rodents and dogs (9). Two human infections appear to stem from bats, one from rabbits and another from cats (*B. koehlerae*). Four genotypes were of uncertain origin – *B. tamiae*, a basal infectious strain that causes febrile illness in humans in Asia and likely originates in rodents (6, 10); *B. bacilliformis*, the causative agent of Carrion’s disease and verruga peruana (11), a South American zoonosis and *B. ancashensis*, a causative agent of verruga peruana (12) and *B. quintana*, the causative agent of trench fever. *Bartonella quintana* has also been found in gerbils (9) and grouped with Old World rodent and bat-associated genotypes, as well as *B. koehlerae* and *B. henselae*, nested within a larger clade of Old World bat-associated *Bartonella*. (Its association with the larger clade of bat-associated *Bartonella* is only evident in the phylogenetic hypothesis based on the 548 bp *gltA* fragment.) Rodent-hosted *Bartonella* has infected carnivores nine times (mostly dogs and cats, but also badgers twice), bats twice and artiodactyls (a cow ectoparasite) once. Bat-hosted *Bartonella* has infected domestic dogs and cats five times and rodents three times. Artiodactyl-associated *Bartonella* has infected a bat and a dog. We also inferred eight transfers of *Bartonella* between rodents and shrews, which are phylogenetically quite distant but presumably share the same terrestrial habitats and some of the same ectoparasite vectors.

Additionally, we noted a minimum of 68 instances in which monophyletic clades or single genotypes contained genotypes isolated from more than one continent/ geographic region, 40 of which spanned both the Old World and New World, denoted in parentheses (Figure S2). Of the clades, 52 (31) involved genotypes found in rodents, 18 (16) involved humans, 20 (15) involved cats and dogs, 2 (1) involved domestic hoof stock, 2 (1) involved badgers, 8 (6) involved shrews, 1 (0) involved pikas and 14 (7) involved bats. Comparing each host category there may be a difference in the likelihood of group carrying global strains of *Bartonella* (Fisher’s exact test, p = 0.1029), and when we grouped humans, cats, dogs and domestic artiodactyls together as human-associated strains and bats, badgers, shrews and pikas together as wildlife-associated strains, the human-associated strains were marginally more likely to be globally distributed (Fisher’s exact test, p = 0.054). All clades known to be found on at least 5 continents were found in humans (*B. clarridgeiae, B. henselae, B. quintana, B. vinsonii* complex and a large clade containing global rodents).

Human-associated strains were also, on average, due to some of the most recent host-shifting events (mean minimum divergence time = 1.6 mya) from one order to another, compared to bats (6.3 mya), carnivores (2.1 mya), shrews (1.1 mya), and rodents (3.0 mya), with the exception of a recent host switch into non-human primates at least ca. 400 kya (Figure S1). As noted above, rodents and shrews seem to be sharing similar *Bartonella* (all shrew-associated host switches are with rodents). If we exclude rodent-shrew transfers, the average host-shift time for rodents is a minimum of 4.8 mya.

## Discussion

### Bartonella *as an environmental bacteria turned insect gut symbiont turned vertebrate pathogen*

The proliferation of studies investigating *Bartonella* in various wildlife populations allows for greater insights into the origins and evolution of *Bartonella* and its potential for spillover more than ever before. Bartonellaceae is nested within the Rhizobiales, a lineage of soil bacteria that contains nitrogen-fixing root-associated members (13). In our study, *B. apis* was the most basal strain of *Bartonella*. Additionally, gut microbiome studies from a variety of insects have revealed that *Bartonella* are actually widespread across arthropods, occurring in carrion beetles, butterflies, bees, various species of ants and a wide variety of ectoparasitic species (14–20). Other studies have hypothesized that perhaps *Bartonella* may have a commensal role in the arthropods that vector it (21, 22). This led us to hypothesize that *Bartonella* originated as an environmental microbe that was picked up by arthropods in which it diversified.

Because most metagenomic studies of bacteria amplify the 16s rRNA gene, there is a large amount of 16s data available and also *Bartonella* can be detected in samples that would not *a priori* be hypothesized to contain *Bartonella*, such as non-hematophagous insects or environmental samples. We mined GenBank for *Bartonella* 16s sequences to test our hypothesis that *Bartonella* is an environmental bacterium that became an insect commensal before becoming a vertebrate pathogen. The 16s rRNA gene is much less powerful for discriminating *Bartonella* species than *gltA* (7) and often metagenomic studies amplify only very small fragments of the gene, making it difficult for us to resolve fine scale diversification but we were able to determine that basal strains of *Bartonella* were largely found in environmental samples and non-hematophagous insects (Figure 1). Additionally, work on *Bartonella* has shown that the evolution of a type 4 secretion system, along with selection on other invasion mechanisms (13), has been instrumental in allowing *Bartonella* to diversify and invade host cells (23, 24) while other work has shown *Bartonella* can incorporate a type 4 secretion system via lateral gene transfer when it coinfects an amoeba with *Rhizobium radiobacter* (25). Further, examinations of lateral gene transfer of metabolic genes in *Bartonella* reveals that many of these genes derive from common insect gut commensal bacteria (26). We strengthen the suggestions of these previous studies by drawing data from insect and environmental metabarcoding studies and demonstrating their basal phylogenetic position within *Bartonella*.

### Bartonella *spillover is predominantly from rodent and domestic animals*

Using the literature (9) and isolates from published sequences on GenBank, we identified 18 genotypes of *Bartonella* that have been found in humans (most of which are also known to cause disease; Table S4). Of these, eight of the genotypes are most closely related to genotypes found in rodents and four are distributed in domestic animals (mostly dogs and cats) but have spilled over into humans. *Bartonella vinsonii* forms a species complex that is associated with transmission to humans from both dogs (subsp. *berkhoffi*) and rodents (subsp. *arupensis*) and we inferred at least 4 separate transfers between humans and these animals based on phylogenetic relationships, however we treated these as a single spillover for the sake of simplicity. Additionally, we identified a genotype of *Bartonella* found in a febrile patient in Thailand (GQ200856) as having over 95% identity with *B. queenslandensis*, a genotype first found in Australian rodents and also found in numerous Asian rodents, suggesting a previously unappreciated rodent-human transmission. Interestingly, one *Bartonella* genotype that was recovered from a Polish forest worker (HM116785) most closely resembled genotypes found in European *Myotis*, a genus of bat, and their ectoparasites (JQ695834, JQ695839, KR822802). That most strains isolated from humans are related to domestic or peridomestic animals strongly indicates that spillover of *Bartonella* requires close contact between humans and the natural reservoirs of these infectious strains.

However, when examined at a broader scale, many of these genotypes lie within larger clades of genotypes found in wild animals. For example, *B. henselae, B. koehlerae* and *B. quintana* were closely related to isolates found in African and European rodent ectoparasites and an Asian bat, nested within a larger clade of Old World bat-associated *Bartonella*. Similarly, *B. mayotiminensis* was closely associated with genotypes of Central American bats detected in this study. This same isolate has also been found in bats in Europe (27) and most recently North America (28). This strongly suggests that bats may be an important reservoir species of potentially zoonotic *Bartonella* strains but that infrequent contact between bats and people prevents transmission. Rather most of the transmission we infer requires the transmission of *Bartonella* into a domestic or peridomestic animal, which can then transmit it to the human. Despite the noted host specificity of *Bartonella* (Table S1), the diversity of strains that infect humans and their distribution across the phylogenetic tree of *Bartonella* suggests that this bacterial genus can and will switch hosts when given the opportunity (especially when hosts are immunocompromised [29, 30]). The relative evolutionary lability of these genotypes is further underscored by the instances in the global phylogeny of genotypes being exchanged between bats and rodents (at least five times).

Overall, we found that rodents were responsible for more transmission of *Bartonella* into humans than any other group, followed by domestic animals. Rodents also transmitted the most *Bartonella* to domestic animals and bats, though infections originating from wildlife such as bats in domestic animals are also relatively common. One potential explanation for the prominence of rodents in host switching may be the generalist tendencies in their ectoparasites. *Bartonella* is vectored by arthropods but some ectoparasites, such as blood sucking hippoboscid flies, are very highly host specific (31) potentially preventing cross-species transmission. In contrast, many rodents host fleas which can bite other taxa and have been found to host many genotypes of *Bartonella* that have originated in rodents and infected other species such as humans (e.g. 32,33). Considerations of the host specificity of the vector species may be very important for determining the risk for disease spillover and indeed public health officials recommend avoidance of potential vectors as the most important measure for prevention of bartonellosis (9).

It is important to note, however, the constraints on our conclusions due to available data. We only have a small fragment of *gltA* to examine across these 860 genotypes, making inferences at deep nodes uncertain. Additionally, we are limited to the animals that have been sampled which are overwhelmingly bats and rodents, as well as symptomatic humans. It is possible and highly likely that there are animal intermediates between these transmission events that are missing.

### *Human movements shape* Bartonella *diversification and infection patterns*

Another interesting pattern that emerged when examining the tree as a whole was the impact of humans in spreading *Bartonella* strains and infections globally. A few particular species that are associated with humans, such as dogs, cats, cows and *Rattus* rats, have managed to bring their strains of *Bartonella* globally (9, 24, 34–39). Rats, in particular the genus *Rattus*, were very common in the largest clade of globally distributed rodent *Bartonella*, with representatives on nearly every continent. This clade also contains four zoonotic genotypes of *Bartonella*, as well as genotypes found in shrews, a dog and a bat ectoparasite, underscoring the important role of human commensals in spreading disease to humans and wildlife and across the globe.

There was a lot of uncertainty in the dating of our divergence times (in one instance two identical genotypes were inferred to be 600,000 years diverged) perhaps due to the small fragment we were able to analyze and the depth of evolutionary history we were exploring. Additionally, there are many genotypes that may have died out or have not been sampled that mean even our minimum divergence date estimates are likely conservative. We cannot therefore state with certainty that humans are responsible for moving other species around, changing disease risk for themselves and wild animals. However, the fact that strains associated with humans or their domestic animals were generally more likely to be found globally, the fact that transfers between humans and domestic animals and other groups were the most recent ones, and the diverse placement of human infections across the phylogeny strongly support a role for humans changing their disease risk as they insert themselves and their associated animals into new habitats and ecosystems.

Such movements and increased human and domestic animal contact with one another and wild animals not only disguises geographic patterns of *Bartonella* diversification (e.g. *B. queenslandensis*, first described in Australia, in *Rattus norvegicus* in California [38]) but it has also led to presumably novel sharing of *Bartonella* between introduced domestic and peridomestic animals and native wildlife. For example, identical genotypes were found in a Chinese *Rattus* individual (DQ986952) and a white-footed mouse, a North American native (AY064534). If the introduced bacteria have adverse fitness consequences, this could be another human-mediated conservation concern.

Other potential aberrant patterns even include transmission of a *Rattus*-associated *Bartonella* into a cow ectoparasite in China (AY517723), the finding of *B. bovis* in an African bat (*E. helvum*, JN172054), a cat (AF071190) and in elk (KB915625) and sharing of *Bartonella* between Old World bats and dogs. The pet trade exacerbates this by shipping exotic animals all over the world, changing the pool of available infections for both the introduced and native species (41). Introduction of domestic species is causing sharing between these species and wild species, changing the disease risks for both.

Overall our findings show that *Bartonella* is a rich system for examining the impacts of humans on patterns of infectious disease spread within species and between species, across landscapes and across the globe. Phylogenetic inferences about the origin of infections should be interpreted with caution as they are heavily influenced by available data and the taxa that have been sampled. There may be many missing links between those we inferred but the hosts simply have not been sampled. Additionally, by analyzing host switching by order we largely overlooked switching between human-associated taxa and wild taxa in the same order (e.g. invasive *Rattus-* associated *Bartonella* transmission to native rodents). At least some part of the noted host specificity of *Bartonella* seems to be due to ecological factors regulating exposure rather than immunological incompatibility. Given the diversity of sources of zoonotic strains, including divergent strains with similar clinical presentations, physicians and researchers should consider a broad range of potential animal hosts and screen for a wide range of *Bartonella* genotypes when investigating the source of a suspected *Bartonella* infection.

## Materials and Methods

In order to ascertain broader patterns of spillover and *Bartonella* transmission between species, sequences were downloaded from Genbank on 30 November 2016 using the search term “Bartonella gltA” and a separate search was conducted using the search term “Bartonella 16s” on 1 February 2017. Insect microbiome studies that detected *Bartonella* were also used (14–20). *Bartonella* from Costa Rican bats in a mosaic agricultural landscape, including previously published (42) and new sequences are also incorporated in this study. Metadata were downloaded from Genbank and/or confirmed by examining the cited publication and are summarized in Supplementary File 1. In the case of data from unpublished work geography was inferred by the host range and/or title information in Genbank. The host of questing ticks was undetermined and therefore denoted as “unknown.” In some cases, genomes of Bartonella strains were published independently from their hosts; in this case we searched other literature to find the source of the strain. Sequences that were not in fact *Bartonella* gltA were removed manually and sequences were aligned using the Geneious alignment algorithm and refined using MUSCLE in Geneious (version 8.1.9 (43). Sequences that were significantly redundant (or multiple sequences of the same species of *Bartonella*) were excluded to reduce the size of the resultant phylogenies. We also excluded some fragments that were too short or had lots of missing data, as well as fragments which misaligned significantly at the ends, causing us to doubt the quality of these end base calls. Alignments were manually inspected and corrected. Two alignments were produced, one of 548 bp and one of 277 bp. The first contained 334 sequences in total and the second included 860 unique sequences.

In order to test for patterns in host specificity and biogeography we also constructed time calibrated Bayesian phylogenies using BEAST 2 (44) for the 548bp fragment and the 277bp fragment. Alignments were split into three partitions based on the base pair’s position in the codon and run in PartitionFinder to determine the best nucleotide substitution models using AICc (45). These parameters were then used to configure the parameters for the BEAST2 run. For the 548 bp run, Partitionfinder determined that all three positions should be run under the same mode, a GTR+I+G+X model; for the 277bp the results were similar except that the model favored was a GTR+I+G. As empirical and maximum likelihood estimated base frequencies usually have little impact on the phylogeny we used observed base frequencies for both sets of nucleotides (45).

We tested three different models for the phylogenetic hypothesis based on the 548bp fragment. All three analyses were run with a gamma site model with empirical base frequencies, an estimated proportion of invariant sites and all nucleotide transition/transversion frequencies except the CT transition rate estimated. The gamma shape prior was set to an exponential distribution with a mean of 1; the proportion of invariant sites was set to a uniform distribution between 0 and 1; all nucleotide substitution rates were set to a gamma distribution with an alpha of 2 and a beta of 0.5 or 0.25 for transitions and transversions respectively. In all cases *Bartonella* was constrained to be monophyletic with *Brucella melitensis* as an outgroup. The first model tested was a strict clock model with a constant population size coalescent model with vague priors as has been used for previous phylogenetic analyses of *Bartonella* (35, 46) with the population size prior set to a 1/X distribution. The second was a birth death model run with a log-normal distributed relaxed molecular clock. The birth rate and relaxed clock mean priors were set to a uniform distribution between 0 and infinity; the relaxed clock standard deviation priors was set to an exponential distribution with a mean of 1; the death rate prior was set to a uniform distribution between 0 and 1. The prior distribution on the divergence date between *Brucella melitensis* and *Bartonella*, the divergence between *Bartonella clarridgieae* and *Bartonella rochalimae, Bartonella coopersplainensis* and *Bartonella rattaustraliani, Bartonella florencae* and *Bartonella birtlesii* were each a normal distribution with a means of 507 mya, 30.8 mya, 82 mya and 57 mya respectively based on previous estimates (47, 48). As we had no prior information about uncertainty of these estimates, we used a standard deviation of 1 my. When setting calibration nodes we only used clades that were well supported and monophyletic in prior analyses of the data regardless of clock model.

The third was identical to the second except that the following calibrations were used: the node at the base of a clade of phyllostomid bat-associated *Bartonella* containing genotypes from three subfamilies (Stenodermatinae, Caroliinae and Glossophaginae) was estimated to have occurred at the divergence of these three subfamilies and was calibrated with a normal distribution with a mean of 24 mya and a standard deviation of 3.76 my based on previous estimates (49–62) collated in TimeTree (48). A nested clade of *Artibeus lituratus* and *Artibeus* watsoni-associated *Bartonella* was estimated to have evolved at their divergence and the prior distribution was estimated with a normal distribution with a mean of 8.5mya (SD = 2.73) based on previous estimates (49–51, 60, 63, 64) collated in TimeTree (48). We used these clades as calibration points as they were strongly supported in all three models, were nested within other Central American bat associated strains and therefore unlikely to have been impacted by human influence and showed evidence of supporting a similar rate of evolution.

For the 277bp tree we also ran PartitionFinder2 (45) which determined that all three positions should be run under the same model, GTR+I+G so we ran our simulations with a GTR distribution, an estimated proportion of invariant sites and a gamma distribution of rates. We tested two models, a strict clock, constant population size coalescent model as described in the first model for the 548bp alignment and a birth death model with a relaxed log normal clock as described in the second model for the 548bp alignment. In both models we constrained *Brucella melitensis* to be an outgroup but no other calibrations were included. All *gltA* model were run for 2.5 × 10^7^ generations and sampled every 50,000 generations.

All *gltA* models converged with all parameters showing an ESS over 200, except a few parameters in the second 548bp model which all showed an ESS over 110. The three models for the 548bp alignment were compared using AICM of the likelihood (65) implemented through Tracer as model comparison using path sampling was not practical. For the 548bp alignment the best model was the third – a relaxed log normal clock calibrated with host divergence dates (dAICM_1st_ = 331.9, dAICM_2nd_ = 21.2). For the 277bp alignment a strict clock was favored over a relaxed clock (dAICM = 153.5). Maximum clade credibility trees were produced using TreeAnnotator, mean heights and a burn in of 10%.

In order to understand the evolutionary origin of *Bartonella* we constructed a phylogeny using sequences from the 16s rRNA gene. All 450 sequences were aligned and trimmed to the same length (259 bp) in Geneious (version 8.1.9; [43]). We constructed a phylogenetic hypothesis in BEAST2 using a strict clock and a birth death model with vague priors as described in the birth death models for the *gltA* genes with *Rhizobium leguminosarum* as an outgroup. The model was run for 10^7^ generations; most ESS were above 300, though the birth rate and death rate ESS were roughly 100. As we were not concerned with speciation dynamics but rather broad topology, we consider this hypothesis to be sufficiently sampled.

This 277 bp *gltA* MCC tree was used in an analysis of host specificity and geographic conservation between related *Bartonella* species. Using the fitDiscrete function in geiger (66), four models of discrete character evolution were fit ‐‐ one using a lambda transformation, one using a white noise transformation, one using an early burst transformation and one using no transformation to model the evolution of host order (with strains isolated from ectoparasites assigned to the ectoparasite’s host) and broad geographic region of isolation both by continent (all except Antarctica) and by Old World versus New World. Fit of the models was assessed using AICc weights and log-likelihoods.

Host switches and sharing of clades between geographic regions was assessed by manually examining the MCC phylogenetic hypothesis based on a 277bp fragment of *gltA*, by examining the location of Genbank records with identical genotypes and by searching the literature for the distribution of named *Bartonella* species. A host switch or geographic shift was inferred so as to capture the minimum number of shifts with posterior support of at least 0.7.

All alignments and metadata are available in the supporting information.

## Acknowledgments

We thank C. Mendenhall, F. Oviedo Brenes, R. Zahawi, W. Figueroa, R. Figueroa, J. Figueroa, Y. Lloria, S. Judson, H. Mao, dozens of Costa Rican landowners, the Organization for Tropical Studies, the Las Cruces Biological Station, and especially J. O’Marr for help with collection of data collection on Costa Rican bat-associated *Bartonella* and Krishna Roskin with obtaining data from Genbank. Additional thanks to Scott Boyd, the Hadly lab, the Boyd lab and Jonathan Flanders for useful comments. This work was graciously funded by the Stanford Woods Institute for the Environment Environmental Ventures Program. HKF was supported by a Bing-Mooney Fellowship in Environmental Science and Conservation and a Stanford Center for Computational, Evolutionary and Human Genomics Postdoctoral Fellowship. Research was approved by the Stanford University Administrative Panel on Laboratory Animal Care (protocol 26920) and conducted under the appropriate Costa Rican permits (RT-044-2015-OT-CONAGEBIO, RT-042-2015-OT-CONAGEBIO, 121-2012-SINAC, RT-019-2013-OT-CONAGEBIO, 226-2012-SINAC).

